# *In silico* identification of novel peptides with antibacterial activity against multidrug resistant *Staphylococcus aureus*

**DOI:** 10.1101/577221

**Authors:** Linda B Oyama, Hamza Olleik, Ana Carolina Nery Teixeira, Matheus M Guidini, James A Pickup, Alan R Cookson, Hannah Vallin, Toby Wilkinson, Denise Bazzolli, Jennifer Richards, Mandy Wootton, Ralf Mikut, Kai Hilpert, Marc Maresca, Josette Perrier, Matthias Hess, Hilario C Mantovani, Narcis Fernandez-Fuentes, Christopher J Creevey, Sharon A Huws

## Abstract

Herein we report the identification and characterisation of two linear antimicrobial peptides (AMPs), HG2 and HG4, with activity against a wide range of multidrug resistant (MDR) bacteria, especially methicillin resistant *Staphylococcus aureus* (MRSA) strains, a highly problematic group of Gram-positive bacteria in the hospital and community environment. To identify the novel AMPs presented here, we employed the classifier model design, a feature extraction method using molecular descriptors for amino acids for the analysis, visualization, and interpretation of AMP activities from a rumen metagenomic dataset. This allowed for the *in silico* discrimination of active and inactive peptides in order to define a small number of promising novel lead AMP test candidates for chemical synthesis and experimental evaluation. *In vitro* data suggest that the chosen AMPs are fast acting, show strong biofilm inhibition and dispersal activity and are efficacious in an *in vivo* model of MRSA USA300 infection, whilst showing little toxicity to human erythrocytes and human primary cell lines *ex vivo*. Observations from biophysical AMP-lipid-interactions and electron microscopy suggest that the newly identified peptides interact with the cell membrane and may be involved in the inhibition of other cellular processes. Amphiphilic conformations associated with membrane disruption are also observed in 3D molecular modelling of the peptides. HG2 and HG4 both preferentially bind to MRSA total lipids rather than with human cell lipids indicating that HG4 may form superior templates for safer therapeutic candidates for MDR bacterial infections.

**Author Summary:** We are losing our ability to treat multidrug resistant (MDR) bacteria, otherwise known as superbugs. This poses a serious global threat to human health as bacteria are increasingly acquiring resistance to antibiotics. There is therefore urgent need to intensify our efforts to develop new safer alternative drug candidates. We emphasise the usefulness of complementing wet-lab and *in silico* techniques for the rapid identification of new drug candidates from environmental samples, especially antimicrobial peptides (AMPs). HG2 and HG4, the AMPs identified in our study show promise as effective therapies for the treatment of methicillin resistant *Staphylococcus aureus* infections both *in vitro* and *in vivo* whilst having little cytotoxicity against human primary cells, a step forward in the fight against MDR infections.

## Introduction

The decline in effective treatment strategies for multidrug resistant (MDR) bacterial infections due to the problem of antibacterial resistance threatens our ability to treat infections now and in the future, and calls for an urgent need to explore new safe drug candidates and alternative treatment strategies^1^. The MDR Gram positive bacteria, methicillin resistant *Staphylococcus aureus* (MRSA), a human opportunistic pathogen, has become a leading causative agent of hospital and community acquired infections over the past few decades, posing a number of challenges for physicians^2–5^. Due to its pathogenicity and potential impact on a large population it is on the World Health Organization (WHO) list of priority pathogens^6^. According to the Centre for Disease Control and Prevention (CDC), MRSA represents a major burden on health care as it can acquire resistance to almost any class of antibiotic^7^, leading to more than 80,000 invasive infections and 11000 deaths each year in the USA alone^8^. The prevalence of MRSA infections in England and Northern Ireland increased for the first time since 2011 from 1.1 in 2016 to 1.3 reports per 100,000 population in 2017^9^. Moreover, treatment of MRSA bacteraemia is a long-standing challenge for the healthcare profession, often complicated by metastatic infections, treatment failure and mortality^4^ Therefore, antimicrobial compounds with new modes of action for the treatment of MRSA infections are urgently needed.

Research into identifying and optimising the use of antimicrobial peptides (AMPs) in infectious disease treatment has been intensifying as they are favoured as a promising new class of therapeutic agents^1, 10^. AMPs have broad spectrum of activity (bacteria, fungi, viruses, parasites etc.), form amphipathic structures, which aid interaction with the cell membrane and have multimodal mechanism of action, which contributes to delayed onset of resistance in host cells against them^11^.

The complex microbial community of the rumen of cattle (*Bos taurus*) adapts to a wide array of dietary feedstuffs and management strategies, and enzymes isolated from this ecosystem have the potential to possess very unique biochemical properties with possible links to economically or environmentally important traits^12^. Indeed, the rumen microbiome has been shown as an underexplored resource for antimicrobial peptide discovery^13–15^ and its potential to contribute some of the much needed urgent alternative therapeutic candidates to tackle the looming problem of difficult to treat multidrug resistant bacterial infections. Advances in nucleic acid-based technology (second generation sequencing, meta ‘omic) and high-throughput sequence analytic methods (advanced bioinformatic approaches) have created new opportunities to investigate the complex relationships and niches within microbial communities, redefining our understanding and improving our ability to describe various microbiomes, including the rumen microbiome. Such an enhanced understanding enable the identification and utilization of the beneficial traits in these microbiomes^16, 17^ Several metagenomic datasets from the rumen have been generated in the last few years, illustrating some of the beneficial traits of the rumen microbiome including the presence of large numbers of novel glycosyl hydrolases^18–27^, esterases^28–30^, lypolytic enzymes^28, 31, 32^, and more recently, antimicrobial compounds^15, 17^, with the latter group of compounds possessing therapeutic potential for the treatment of multi-drug resistant bacteria.

Here we combined the application of metagenomics, using one of the largest rumen metagenomic dataset^19^ available, with advanced computational analytic tools and chemical models to identify and characterise AMP candidates for the treatment of MDR infections. This metagenomic data set contains more than 268 Gb, or 1.5 billion read pairs, of metagenomic DNA from one single sample with the DNA from microorganisms that colonized plant fiber during incubation in the cow’s rumen. *De novo* assembly of reads resulted in more than 2.5 million predicted open reading frames at an average of 542 bp and 55% predicted full-length genes. We employed the classifier model design, a feature extraction method using molecular descriptors for amino acids for the analysis, visualization, and interpretation of AMP activities, and for the *in silico* discrimination of active and inactive peptides in order to define a small number of promising new lead AMP test candidates for chemical synthesis and experimental evaluation from this dataset. We also show the innocuity *ex vivo*, and the anti-MRSA efficacy of two of these AMPs both *in vitro* and *in vivo*.

## Results and Discussion

### *In silico* prediction and identification of AMPs using computational analysis

Following, the first selection criteria in the computational analysis, 917,636 sequences (36%) of the 2,547,270 predicted protein sequences in the Library “Cow” remained (i.e., protein sequences with a maximum length of 200 amino acids (AAs)^33^ and not more than 5% unknown AAs (marked by X, *). Of these 917, 636 sequences, only 829 sequences fulfilled the criteria of AA distances (AAD) <0.2 or a small AA pair distances (AAPD) <1.45, ensuring their potential to be AMPs. For example, only 65 sequences met these criteria in the first 68, 274 sequences analysed, with isolated points outside a relatively dense distribution area as illustrated in Fig. 1a. Descriptor computations generated positively charged loading - hydrophobicity plots, indicating that the selected sequences from the Library “Cow” are represented in only a small portion of all AMP regions from the Library “AMP” (see Fig. 1b). Results from each computational step used in the identification of novel AMPs from the Hess et al^19^ rumen metagenomic dataset is summarized in the supporting information (SI) Table S1.

**Fig. 1.**
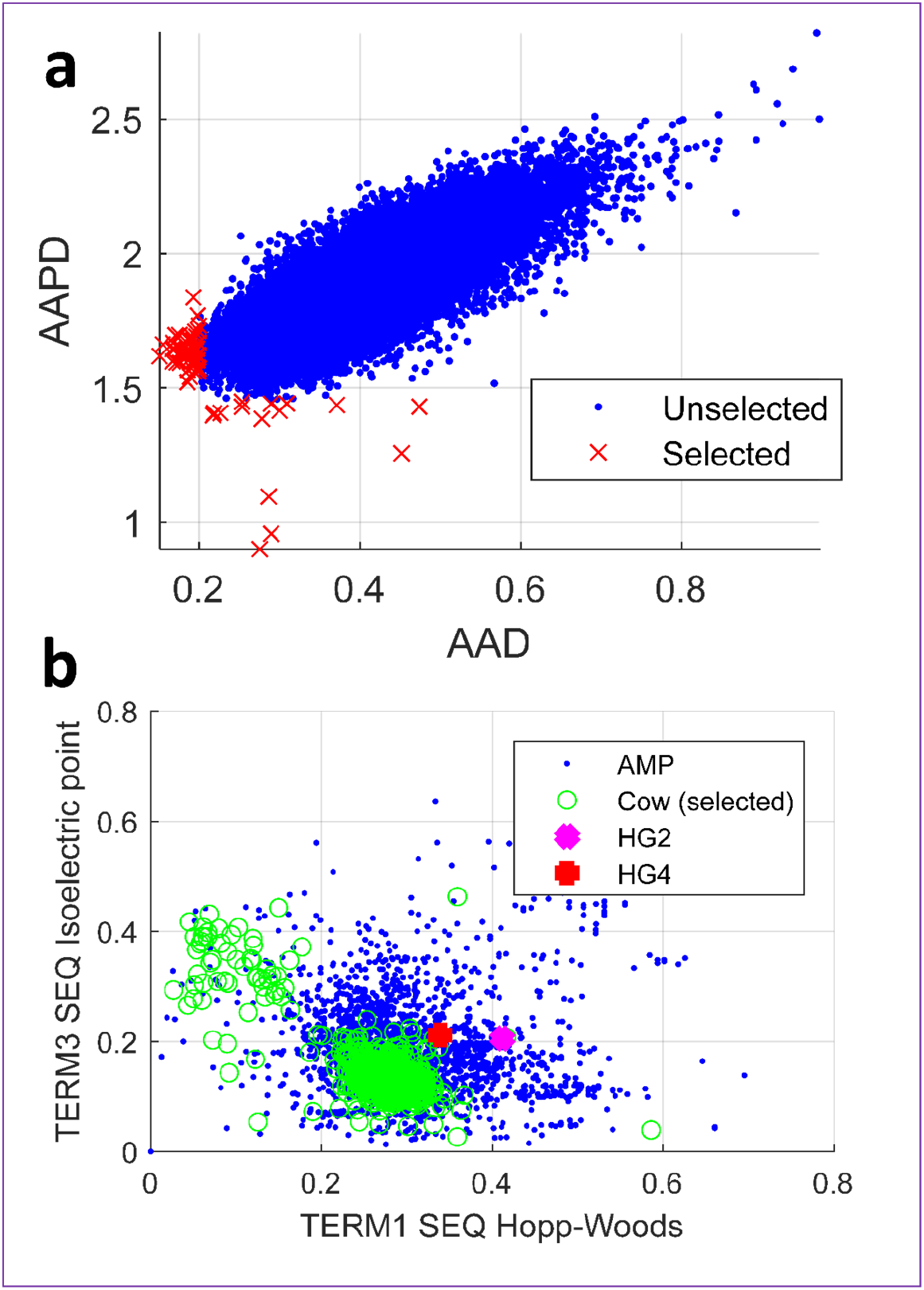
**a)** visualization of distances for AA acids (AAD) and AA pairs (AAPD) for the first 68,274 sequences from library “Cow”^19^ meeting the first selection criteria: candidates with AAD<0.2 or AAPD<1.45 are selected as candidates (here: 65). **b)** standard hydrophobicity (TERM1 SEQ Hopp-Woods) - loading (positively charged, TERM3 SEQ Isoelectric Point) plot. Blue dots are known AMPs (library “AMP” consisting of AMPs from the APD2^34^ and Hilpert Library^35^), green colored signs are AMP hits identified from library “Cow”, and finally selected peptides HG2 (magenta) and HG4 (red).

Six most promising AMP sequences (termed Hess-Gene 1-6 (HG1-HG6) (SI Table S2) were identified from the 829 sequences we utilized *in silico* approaches (Materials and Methods) sequences. The corresponding nucleotide sequence and additional information for the identified genes can be found in supporting information S3.

Two of these six promising candidates, HG2 (MKKLLLILFCLALALAGCKKAP) and HG4 (VLGLALIVGGALLIKKKQAKS) containing 22 and 21 AA residues respectively, were selected for subsequent characterisation. Sequence homology analysis using NCBI’s BLASTP^36^ against the non-redundant (nr) protein sequences suggests that HG2 is most similar to hypothetical or uncharacterised proteins with unknown functions from *Treponema maltophilum* ATCC 51939, *Pedobacter soli* and *Bacteroides* sp., while HG4 is most similar to Na^+^/H^+^ antiporter NhaC family protein from *Fibrobacter* sp. UWR2, and a hypothetical protein from *Bifidobacterium adolescentis* (Table 1).

**Table 1.** Homology of antimicrobial peptides HG2 and HG4 to known sequences

| Assigned AMP name |  | HG2 | HG4 |
| --- | --- | --- | --- |
| Sequence |  | MKKLLLILFCLALALAGCKKAP* | VLGLALIVGGALLIKKKQAKS* |
| Amino acid (AA) (length) |  | 22 | 21 |
| Location on cow dataset |  | NODE_664976_length_19740_cov_2.033485_orf_00810_19724..19789 | NODE_3958153_length_85376_cov_8.525382_orf_203250_82784..82849 |
| Most similar homolog on APD2 (stop codon '**' removed) | APD ID | AP00494<sup>37</sup> | AP01737<sup>38</sup> |
|  | Similarity % | 40% | 48% |
| Most similar homolog on NCBI blastp (stop codon '**' removed) | Accession number | EPF31931.1 | WP_088636977.1 |
|  | Description | Hypothetical protein HMPREF9194_02286 [*Treponema maltophilum* ATCC 51939] | Na<sup>+</sup>/H<sup>+</sup> antiporter NhaC family protein [*Fibrobacter* sp. UWR2] |
|  | Score bits/Identities %/<br>E-value | 43.5 bits (95)/ 17/24(71%) / 0.001 | 44.8 bits (98)/ 16/19(84%)/ 0.001 |

Similarly, homology analysis of the AMP sequences against the proteins from WGS metagenomic projects (env_nr) suggests that HG2 is most similar to hypothetical proteins from marine metagenomes, while HG4 is most similar to permease of the drug/metabolite transporter (dmt) superfamily identified from a hydrocarbon rich ditch metagenome and hypothetical proteins from marine metagenomes. It is important to note that the AMPs match to only a small part of their homologous sequences. The high e-values and low bit scores and coverage of the homologous sequences to HG2 and HG4 (as the best hits still had relatively high e-values of 0.001), as well as the fact that they are automatically curated unreviewed sequences should also be noted, as this indicates the potential novelty of the peptides. Therefore, this study has not only identified HG2 and HG4 sequences from the rumen metagenomic dataset as antimicrobial proteins but may also provide insight into the function of the full proteins carrying their homologous sequences currently annotated as hypothetical or uncharacterised proteins for the first time.

HG2 and HG4 were chemically synthesized as linear, C-terminal amidated peptides on resin (≥95% purity, see SI Fig. S1 for mass spectrometry analysis and peptide synthesis reports) using solid phase Fmoc peptide chemistry^39^ before their antimicrobial activity was investigated. It should be noted that peptide HG2 was synthesised with a disulphide bond linking cysteine residues at positions 10 and 18, since HG2 showed little antimicrobial activity when lacking this disulphide bond (results not shown). Similar to previously reported antimicrobial peptides^40–42^, HG2 (C_111_H_196_N_26_O_23_S_3;_ MW=2359.12 Da) and HG4 (C_99_H_182_N_26_O_24;_ MW=2120.69 Da) are cationic both having a net positive charge of +4. Whereas a hydrophobicity ratio of 57% was calculated for HG4 using ExPASy’s ProtParam tool^43^, a hydrophobicity ratio of 72% was predicted for HG2, which is unusually high compared to the ratio that has been reported for most AMPs^44–47^. This puts HG2 into the small group of AMPs, representing <1% of AMPs deposited in the APD3 database^33^, for which a hydrophobicity ratio of ≥72%^33^ has been reported. The positive charge and hydrophobicity of AMPs are known to contribute of their antimicrobial activity as they play a role in their ability to interact with the bacterial cell membrane^48^.

### 3-Dimensional molecular modelling of peptide structures

Three dimensional structural modelling of HG2 and HG4 using PEPFOLD^49^ suggests that these peptides have a high proportion of helical content (Fig. 2). In the case of HG2, the cysteine bond stabilises the capping of the helix and the C-terminus region. Noteworthy is the clear amphipathic nature of the helix with the hydrophobic residues, particularly, Leu, aligned in a typical Leu-zipper motif. The Nt-C-termini includes a high proportion of the charge residues (Lys), which also contribute to the segregation of charges along the peptide. As shown for other examples^50^, the amphipathicity of peptides, and in particular, peptides with helical conformation is an important feature of antimicrobial peptides that explains their membrane disruptive mechanism of action.

**Fig. 2.**
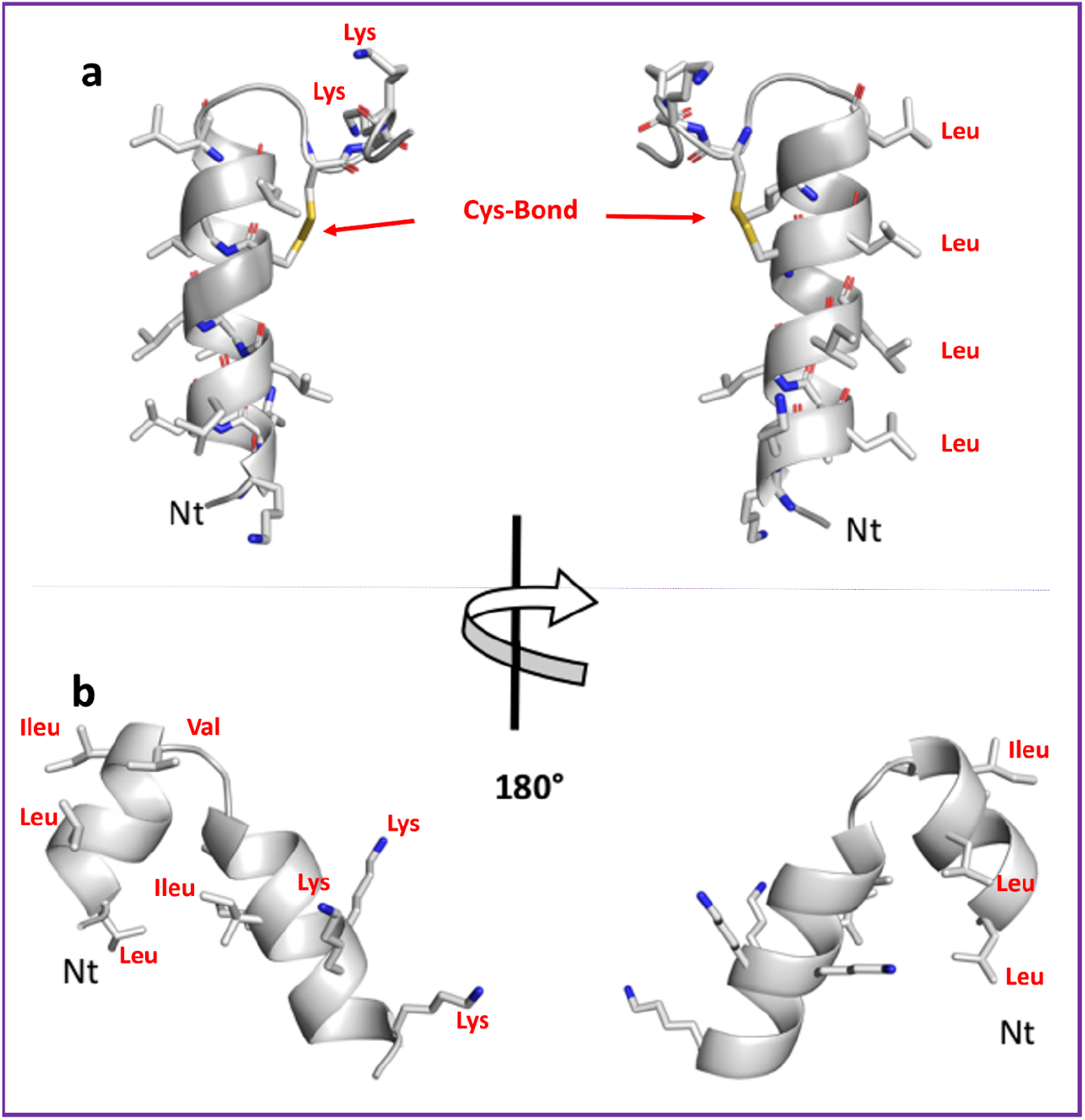
Predicted 3D structures for peptides: **a)** HG2, **b)** HG4. Main-chain and side chains depicted in ribbon and stick representation respectively and coloured according to atom type: Carbon, Oxygen and Nitrogen in green, red and blue respective. Two orientations are shown rotated about the shown axis. Ct and Nt as well as selected residues are depicted in the figure. Figures were rendered using PyMol.

Structural modelling also shows a high content of helical conformation in HG4 (Fig. 2). The peptide forms a helix-turn-helix motif with the C-terminal helix capping stabilised by hydrophobic interactions between the helices. The distribution of charges is asymmetrical as expected, given the sequence of the peptide with the C-terminal part including all charged residues (Lys mainly). The N-terminal helix contain mainly hydrophobic residues and the C-terminal helix all charged residues with the exception of the first turn of the helix, containing a high proportion of hydrophobic residues that form a mini core with the previous helix possibly stabilising the conformation of the motif. The resulting conformation of the peptide is therefore an amphipathic molecule, albeit different from HG2, could also point to a mechanism of action on membranes.

### Antimicrobial susceptibility studies

#### Determination of Minimum inhibitory concentrations (MIC)

We determined the antibacterial activity of HG2 and HG4 against various clinically important multidrug-resistant pathogens including strains of *Acinetobacter baumannii*, *Klebsiella pneumoniae, Pseudomonas aeruginosa, Escherichia coli*, *Salmonella enterica* serovar Typhimurium, *Staphylococcus aureus, Bacillus cereus, Enterococcus faecalis* and *Listeria monocytogenes* (Table 2). HG2 and HG4 had favourable antibacterial activity mostly against Gram-positive pathogens, including multidrug resistant (MDR) strains (Table 2), and were most potent against methicillin resistant *Staphylococcus aureus* (MRSA) strains. HG2 had a minimum inhibitory concentration (MIC) range of 16-32 μg/ml while HG4’s MIC was 32-64 μg/ml depending on MRSA strain, falling within the range of MICs for other rumen-derived AMPs we identified previously^15^, and for commercially available AMPs from isolates^51, 52^. The peptides also showed activity against some Gram-negative bacteria strains, specifically some non-resistant *A. baumannii* strains and *P. aeruginosa* strains C3719 and LES400 isolated from cystic fibrosis patients (Table 2).

**Table 2:**
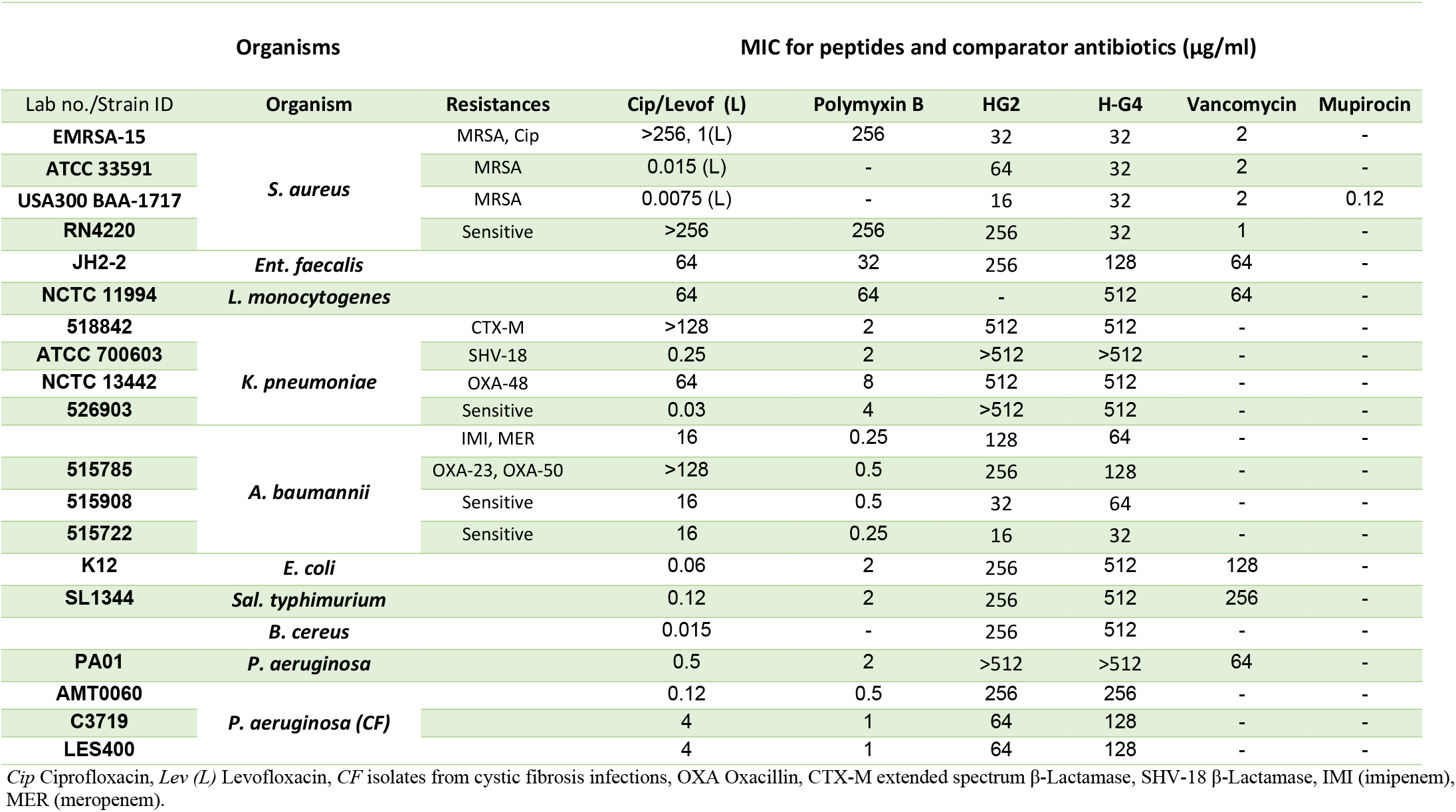
MDR bacteria susceptibility to HG2, HG4 and comparator antibiotics measured by MIC

#### Time kill kinetics

The bactericidal activity of HG2 and HG4 against logarithmic-phase MRSA USA300 cells was investigated by time kill kinetic studies. Compared with vancomycin and mupirocin, HG2 and HG4 (at 3x MIC concentration) had a rapid bactericidal activity against MRSA USA300 strain (Fig. 3a), causing reductions of >3 log_10_ CFU/ml and >6 log_10_ CFU/ml, respectively, within the first 10 min. HG2 and HG4 induced complete cell death within 5 hours and 10 min of treatment respectively, with no recovery observed after 24 hours of incubation. This rapid and total loss in bacteria cell viability is similar to the killing kinetics that have been reported for many fast acting antimicrobial peptides^15, 53^. As expected, vancomycin and mupirocin at 3x MIC produced ≥2 log_10_ CFU/ml reductions attributable to differences in kill kinetics and mode of action^54^.

**Fig. 3.**
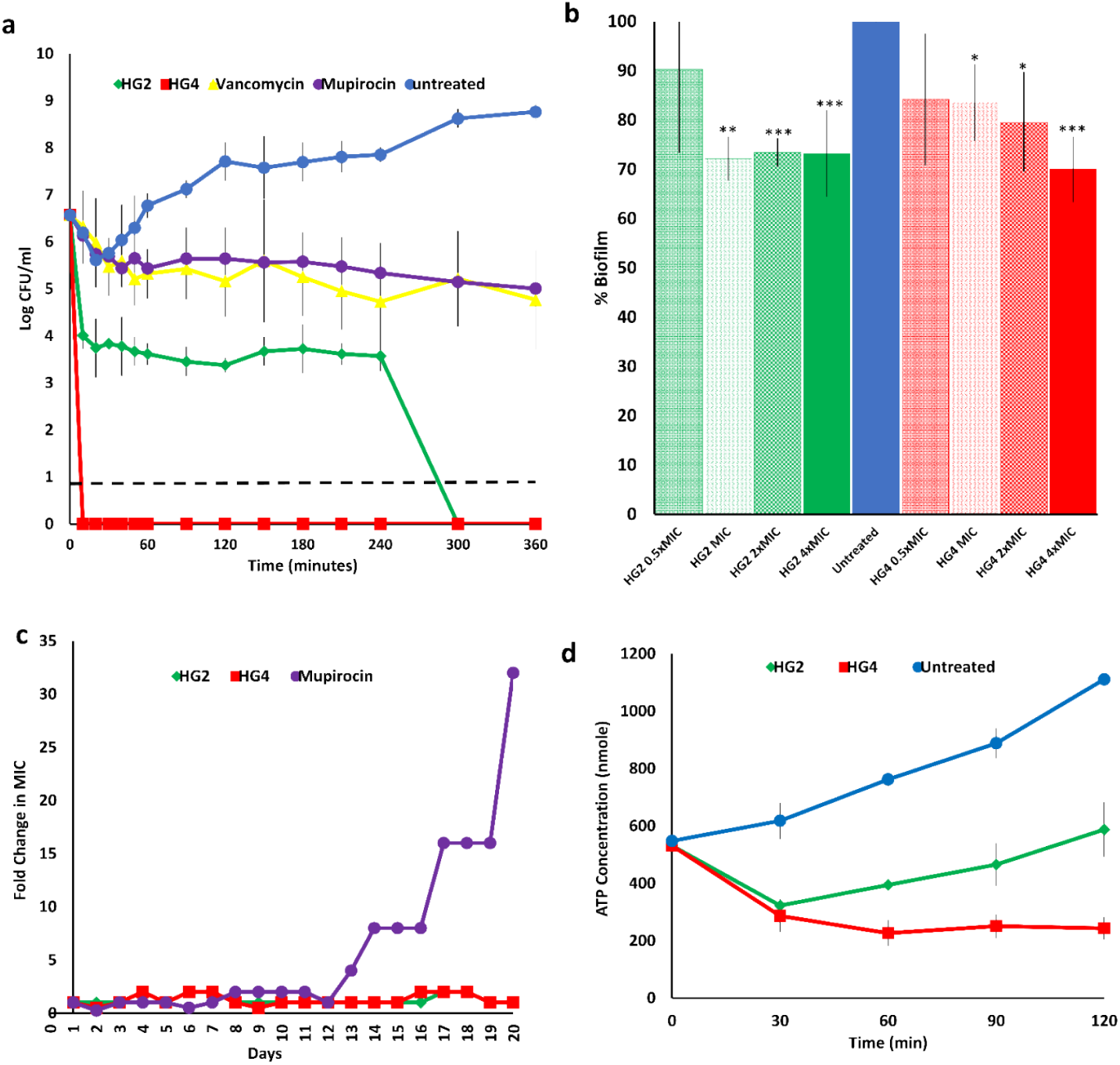
Antimicrobial susceptibility and activity of HG2 and HG4. **a)** Time dependent kill of MRSA USA300 cells by AMPs at 3x MIC concentration. Dashed lines indicate limit of detection. **b)** Anti-biofilm activity against MRSA USA300 biofilms: *, ** and *** (P ≤ 0.05, 0.01 and 0.001 respectively-significantly different from untreated cells (positive). **c)** Resistance acquisition during serial passaging of MRSA USA300 cells in the presence of sub-MIC levels of antimicrobials. The y axis is the fold change in MIC during passaging. For mupirocin, 32x MIC was the highest concentration tested. The figure is representative of 3 independent experiments. **d)** ATP depletion activity in MRSA USA300 cells.

#### Anti-biofilm activity

There have been numerous AMPs that have been reported to be capable of inhibiting biofilm formation for difficult to treat pathogens, favouring their application as antimicrobial agents in medical implants and other biomaterials^55–61^. This prompted us to utilize a 96-well biofilm model^15^ to investigate the ability of HG2 and HG4 to dislodge/disrupt and disperse already formed and established MRSA USA300 biofilms. In general, all AMP treatments showed activity against established biofilms at MIC, 2x MIC and 4x MIC concentrations. No statistically significant anti-biofilm activity was observed in biofilms treated with 0.5x MIC AMP concentrations (Fig. 3b). The anti-biofilm activities of HG2 and HG4 indicate their suitability potential agents for the disinfection of medical devices as well as in the treatment of biofilm infections such as wounds.

#### Selection for resistance (serial passage)

Although relatively uncommon, bacterial resistance to cationic antimicrobial peptides is an evolving phenomenon^62, 63^. Resistance to many AMPs including polymyxin B has recently been reported^64–66^, and it is therefore important to understand bacterial resistance to AMPs and to identify and design more robust AMPs. Mechanisms of resistance to AMPs, which are mostly non-specific and confer moderate levels of resistance^67, 68^, are mainly based on changes in the physicochemical properties of surface molecules and the cytoplasmic membrane^62, 63^. For therapeutic AMP candidates, it is important that bacterial AMP resistance, which may develop due to selective pressure is not based on mutations or acquisition of specific resistance genes, which can then be horizontally transferred between bacteria species as with conventional antibiotics^69, 70^. Here, we assessed the likelihood of resistant mutants and/or resistance arising when MRSA cells are exposed to sub-MIC levels of HG2 and HG4. Continuous exposure of bacteria cells to sub-lethal doses of the AMPs over a period of 20 days did not produce resistant mutants (Fig. 3c), and MICs remained within 1-2 fold increases compared to mupirocin treated cells, which had a 32-fold MIC increase within the same period. The observed increase in MIC is rather common for many AMP-based molecules as a small change in the MIC after exposure to the AMP is to be expected^71, 72^. Our inability to recover resistant mutants in this experiment suggests that the HG2 and HG4 may have non-specific or multiple cellular targets as has been described previously for peptides^73^.

### Biochemical mode of action studies

#### ATP depletion assay

Adenosine triphosphate (ATP) is a high-energy nucleoside triphosphate molecule formed in the cytosol of bacteria and mitochondria of eukaryotes and drives most cellular and metabolic processes in microbial cells^74–76^. Changes in the concentration of ATP can be used as an indicator of cell viability and competence. We tested the effect of HG2 and HG4 on ATP concentration levels in *S. aureus* MRSA USA300. As expected, untreated bacteria cells generated increasing amounts of ATP over time, whereas significantly lower concentrations of ATP were observed in HG2 and HG4 treated cells (Fig. 3d). This decrease in ATP concentrations may be an indication of ATP depletion, limiting cellular energy and thus other related cellular processes (such as substrate transport, homeostasis and anabolism) likely linked to cell membrane disturbance. HG2 and HG4 induced a significant (*P* = 0.018 and 0.003, respectively) decrease in ATP concentration in *S. aureus* MRSA USA300 cells, which is similar to what was reported for other cationic AMPs^47, 77^ Hilpert et al^47^ demonstrated that many short AMPs had a strong effect on ATP concentration whereas several 26mer α-helical peptides did not.

#### Bacterial membrane permeabilisation assay

Since HG2 and HG4 peptides possessed rapid bactericidal effect, we used the propidium iodide (PI) method^15, 78^ to determine if these AMPs were able to permeabilise MRSA USA300 cytoplasmic membrane similar to what was observed for other rumen derived AMPs^15^. MRSA USA300 cells exposed to increasing concentrations of HG2 or HG4 both showed increase in PI entry/fluorescence over time, demonstrating that they were able to permeate the cytoplasmic membrane and therefore indicating that they may possess pore-forming activity (Fig. 4a, b). Indeed, significant permeabilisation (p < 0.01) of MRSA USA300 cytoplasmic membrane was observed even at sub-MIC concentrations, e.g. as low as 1 and 16 μg/ml for HG2 and HG4, respectively (MIC values being 16 and 32 μg/ml for HG2 and HG4 respectively). The Effective Concentration 50 (EC50) (defined as the concentration of a drug at which the drug is half-maximally effective) of HG2 and HG4 measured after 80 min of incubation were 1.351 (±0.27) and 13.85 (±3.22) μg/ml, while total membrane permeabilisation was observed at 3.9 and 62.4 μg/ml, respectively (Fig. 4c). Membrane permeabilisation kinetics of HG2 and HG4 at their MIC concentration, showed that HG2 was able to permeabilise the membrane faster (80% permeabilisation at 1 min and maximal effect after 5 min) than HG4 (minor permeabilisation at 20 min, maximal effect after 40 min) (Fig. 4d).

**Fig. 4.**
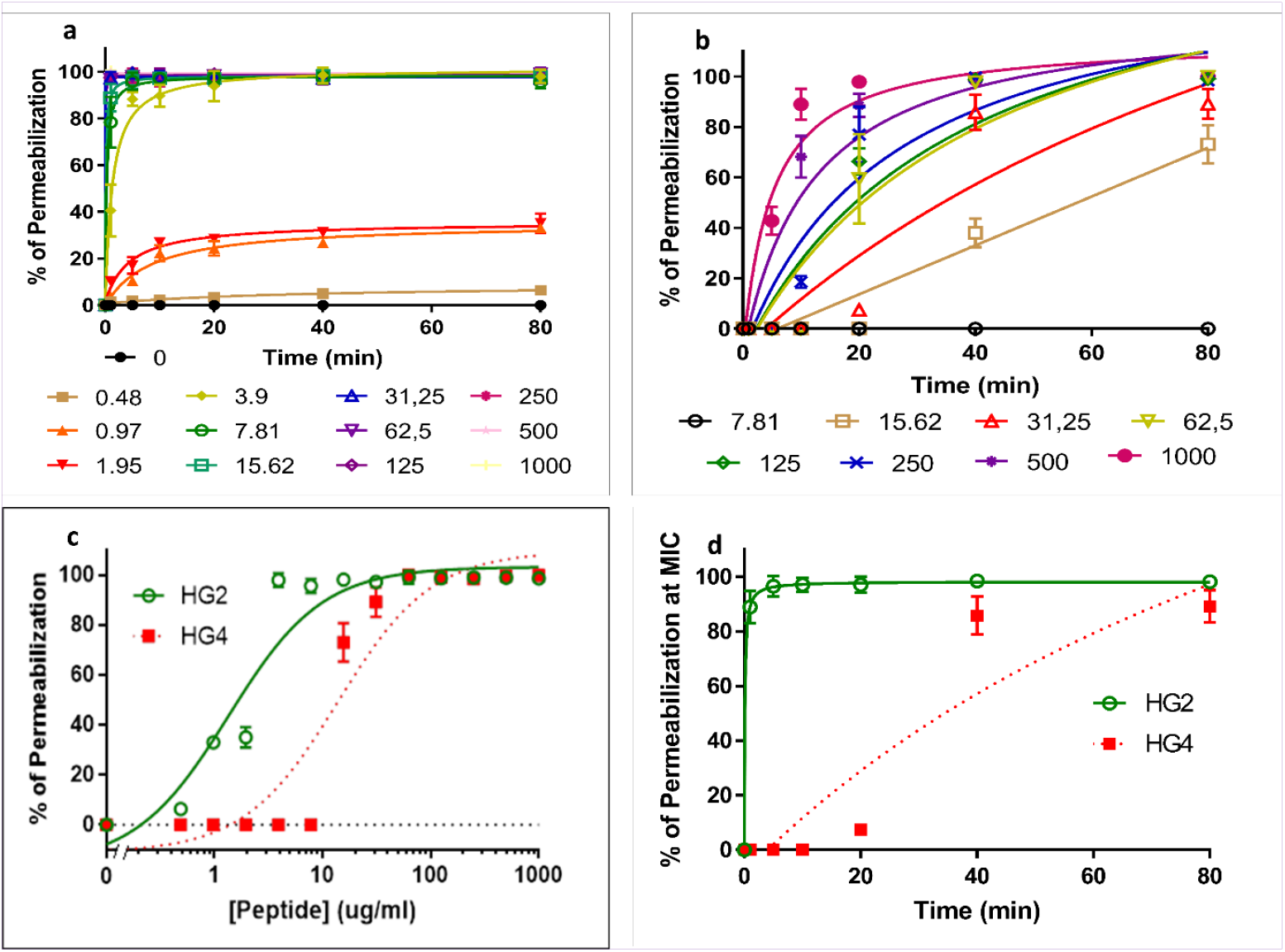
Membrane permeabilisation action of HG2 and HG4 against MRSA: **a)** Membrane permeabilization activity of HG2 at different concentrations (μg/ml) against MRSA USA300 cells measured by propidium iodide assay over time. **b)** Membrane permeabilization activity of HG4 at different concentrations (μg/ml) against MRSA USA300 cells measured by propidium iodide assay over time. **c)** Determination of EC50 (Effective Concentration 50) of HG2 and HG4 membrane permeabilisation measured after 80 min. **d)** Membrane permeabilisation kinetics of HG2 and HG4 at their MIC concentration. In all cases, values are from three independent replicates; results are expressed as means ± standard deviation).

#### Transmission electron microscopy

Transmission electron micrographs (TEM) of cells treated with HG2 and HG4 at 3x MICs for 1 h revealed changes in cell morphology and some cytoplasmic damage (Fig. 5). The morphological changes observed in the HG2 and HG4 treated MRSA USA300 cells correspond with the membrane permeabilisation activity of the peptides. The semi-quantitative nature of TEM analysis means that investigation into events leading up to changes in cell morphology may be necessary.

**Fig. 5.**
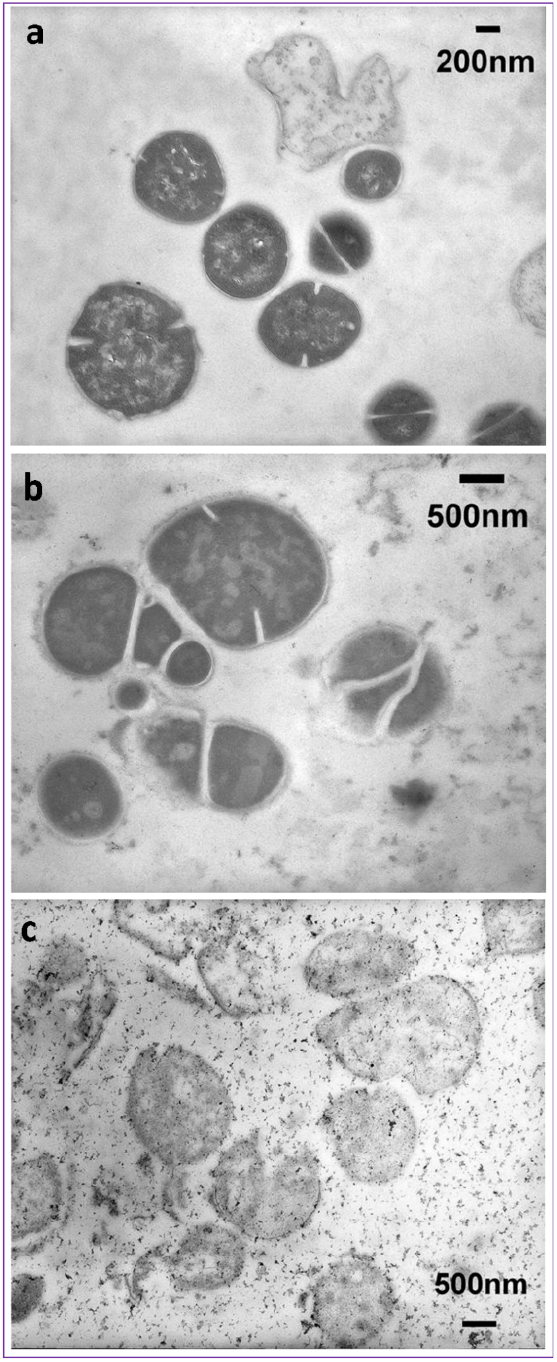
Representative transmission electron micrographs of MRSA cells. **a)** micrographs untreated MRSA USA300 cells. **b)** HG2 treated (3x MIC for 1 h) MRSA USA300 cells. **c)** HG4 treated (3x MIC for 1 h) MRSA USA300 cells. Scale bars are 200 or 500 nm as shown on micrographs.

### *In vitro* and *ex vivo* innocuity and cytotoxicity studies

#### Haemolytic activity

To establish the potential of HG2 and HG4 as therapeutic agents, the haemolytic effect of HG2 and HG4 were tested on human red blood cells. HG2 and HG4 induced low haemolysis, with HC_50_ (i.e. the concentration of peptide causing 50 % haemolysis) of 409 ±67 and 458 ±101 μg/ml, respectively, with safety factors of 26.2 and 14.6X MIC respectively (Table 3).

**Table 3.** Cytotoxicity and haemolytic activities of HG2 and HG4 on human cells. Cytotoxicity is expressed as IC_50_ (i.e. the concentration of peptide in μg/ml causing a reduction of 50% of the cell viability). Haemolytic activity is expressed as HC_50_ (i.e. the concentration of peptide causing 50% haemolysis). IC_50_ and HC_50_ are expressed in μg/ml concentrations. Therapeutic Indexes (T.I) corresponding to the fold difference between IC_50_ or HC_50_ and MIC values (for MRSA USA300) are given in brackets.

| Cell type (human) | HG2 (X MIC) | HG4 (X MIC) |
| --- | --- | --- |
| **BEAS-2B** | 120 +/- 25 (X 7.7) | >1000 (> 32) |
| **HEPG2** | 359 +/- 76 (X 23) | >1000 (> 32) |
| **IMR-90** | 96 +/- 21 (X 6.1) | 294 +/- 42 (X 9.4) |
| **Erythrocytes** | 409 +/- 67 (X 26.2) | 458 +/- 101 (X 14.6) |

#### Cytotoxicity studies on human primary cells and cells lines

Cytotoxicity of HG2 and HG4 against different human cell types was evaluated by measuring the IC_50_, which is the concentration of peptide inhibiting 50% of the cell viability. Lung fibroblast (IMR-90) cells were found to be the most sensitive to HG2 and HG4 with IC_50_ of 96 ± 21 and 294 ± 42 μg/ml for HG2 and HG4. Lung epithelial (BEAS-2B) and liver (HepG2) cells) cells were the least sensitive to HG2 and HG4 with IC_50_ of 120 ± 25 and 359 ± 76 μg/ml for HG2 and IC_50_ >1000 μg/ml for HG4 (Table 3). As a whole, cytotoxicity data showed that HG4 was less toxic than HG2 for human cells and that epithelial cell types (BEAS-2B and HEPG2) were the least sensitive, while fibroblast and endothelial cells were more susceptible to the peptides (Table 3). The high hydrophobicity of HG2 may contribute to its higher toxicity to human erythrocytes and cell lines. This is similar to Gramicidins that possess high hydrophobicity ratios and are exclusively used topically due to their haemolytic side-effects^79–81^, it is possible that the future application of HG2 might be restricted to topical application to treat superficial infections, unless modified derivatives/analogues of HG2 with improved cytotoxicity become available.

#### Peptide-lipid interaction and insertion assay

The interaction of HG2 and HG4 with lipid monolayers was measured by the critical pressure of insertion, reflecting the affinity of the peptides for specific lipids. Insertion capacity was first measured using total lipids extracts obtained from MRSA USA300 cells or human erythrocytes (Fig. 6a, b, and SI Table S4) and obtained data suggested that HG2 and HG4 had higher affinity and insertion ability in MRSA lipids, with critical pressure of insertion of 35.07 and 30.99 mN/m and 42.59 and 44.18 mN/m for HG2 and HG4 in MRSA and erythrocyte lipids, respectively. These results showing that HG4 is less able than HG2 to insert into erythrocyte lipids is in accordance with the lower haemolytic activity observed in HG4 compared to HG2 (HC50 of 409 and 458 μg/ml for HG2 and HG4, respectively).

**Fig. 6.**
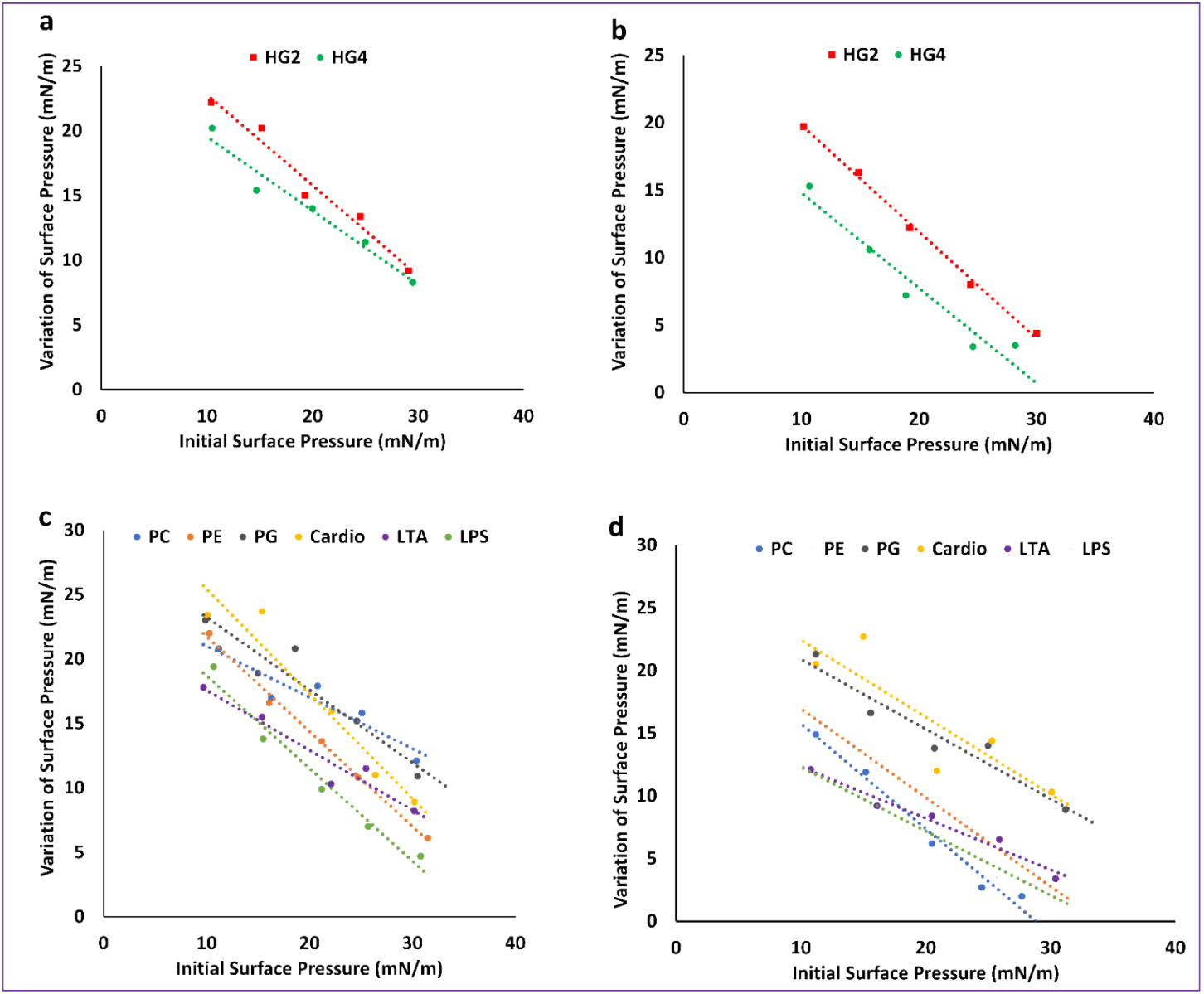
Peptide lipid interaction and insertion measurements: **a)** Interaction of HG2 and HG4 (at 1 *μ*g/mL final concentration) with lipids (either total lipid extracts or pure lipids) was measured using lipid monolayers. a) interaction HG2 and HG4 with total MRSA lipid extract. **b)** interaction HG2 and HG4 with total lipid extract from human erythrocytes. **c)** interaction of HG2 with pure lipids and c) interaction of HG4 with pure lipids. 1-palmitoyl-2-oleoyl-sn-glycero-3-phospho-(1’-rac-glycerol) (PG), 1-palmitoyl-2-oleoyl-sn-glycero-3-phosphoethanolamine (PE), Cardiolipin (Cardio), Lipoteichoic acid (LTA) from *S. aureus*, Lipopolysaccharide (LPS) from *E. coli* and (1-palmitoyl-2-oleoyl-glycero-3-phosphocholine (PC).

To identify the lipid partner(s) recognized by HG2 and HG4 in bacterial and eukaryotic membranes measurement of the critical pressure of insertion in pure lipids were performed (Fig. 6c, d and SI Table S4). Results indicated that HG2 and HG4 had different affinities and insertion capacities in pure lipids from bacteria or eukaryotes. Whereas HG4 interacts preferentially with bacterial lipids expressed in the outer leaflet of the membrane (1-palmitoyl-2-oleoyl-sn-glycero-3-phospho-(1’-rac-glycerol) (POPG or PG), 1-palmitoyl-2-oleoyl-sn-glycero-3-phosphoethanolamine (POPE or PE), cardiolipin, lipoteichoic acid from *S. aureus* (LTA), lipopolysaccharide from *E. coli* (LPS) over eukaryotic lipids (1-palmitoyl-2-oleoyl-glycero-3-phosphocholine POPC or PC) (i.e. PG > cardiolipin > LTA > LPS > PE > PC), HG2 displayed the opposite order of selectivity, with first the major lipid present in the outer leaflet of the eukaryotic membrane (PC) followed by bacterial membrane lipids (i.e. PC > PG > LTA > Cardio > PE > LPS). These observations are in accordance with the higher toxicity of HG2 (high susceptibility of human cells to HG2) against human cell lines compared to HG4 (Fig. 6c, d and SI Table S4). Interestingly, neither HG2 nor HG4 insert efficiently into LPS. This corresponds to the low antimicrobial activity of HG2 and HG4 against Gram negative bacteria compared to their potent activity against Gram positive bacteria). Measurement of the speed of insertion of HG2 and HG4 in total lipid extracts or pure lipids at an initial surface pressure of 30 ± 0.5 mN/m (corresponding to the theoretical surface pressure of eukaryotic and prokaryotic membranes) showed that HG2 inserts faster into all lipid monolayers compared to HG4 (Table 2 in supporting information), confirming results obtained in the membrane permeabilisation assay, which indicated a faster bacterial membrane permeabilisation of HG2 when compared to HG4.

#### *In vivo* efficacy studies

*Galleria mellonella* has been used widely as an effective model for testing antimicrobials drugs and their toxicity^82–85^. We investigated the toxicity of HG2 and HG4 against *G. mellonella* larvae as well as their ability to protect the larvae from a lethal dose of an MRSA USA300 infection. The peptides HG2 and HG4 showed no toxic effect in *G. mellonella* at either 1x MIC or 3x MIC concentrations (Fig. 7a). As with the control group of larvae, no phenotypic modification such as melanisation and/or motility reduction was observed in larvae injected with HG2 and HG4.

**Fig. 7.**
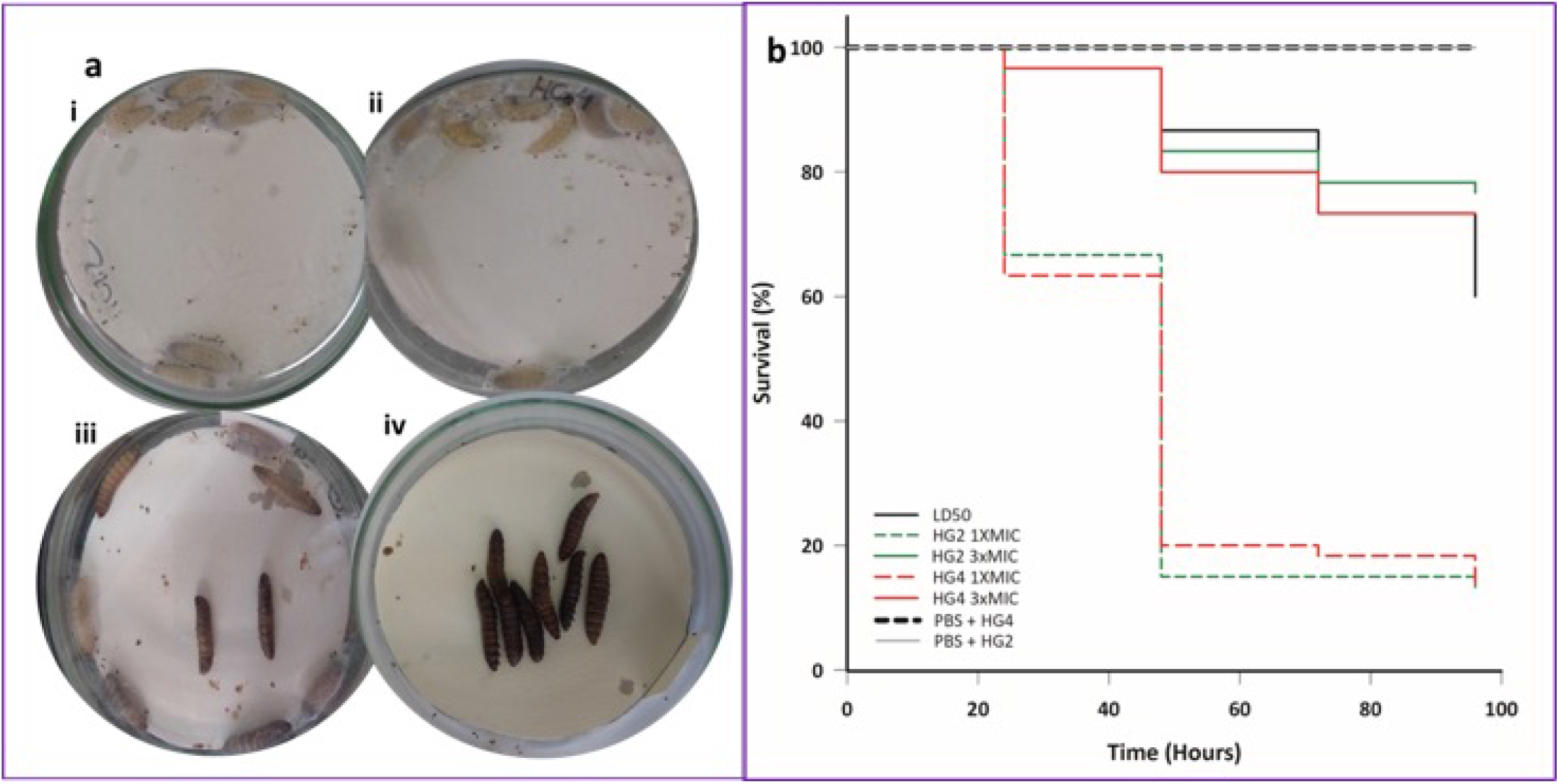
*In vivo* efficacy assessment in *G. mellonella* MRSA infection model: **a)** representative images of toxicity assay of peptides-**i)** HG2, and **ii)** HG4 in *G. mellonella* 120 h post treatment with 3x MIC concentrations. The larvae remained alive and without melanisation. **iii)** virulence assay of MRSA USA300 in *G. mellonella* using a lethal dose inoculum of 10^6^ CFU/per larvae-iii) 24 hours post infection: some larvae were dead and partial melanisation was observed. iv). 48 hours post infection: most larvae were dead and complete melanisation was evident. The experiment was done with three experimental replicates, each containing groups of 10 larvae. **b)** Kaplan-Meier survival curves of *G. mellonella* infected with a lethal dose of *S. aureus* (2.25 x 10^6^ CFU/larvae) and treated with placebo (showing a 100% larvae survival rate) or peptides HG2 and HG4 at a 1x and 3x MIC concentrations.

The MRSA USA300 infective dose (LD_50_) was determined as being 10^5^ CFU/larvae, while the lethal dose (LD) was determined as 2.25×10^6^ CFU/larvae (caused melanisation and death of all larvae within 24 h (Fig. 7a). Larvae infected with MRSA USA300 LD and treated with peptides HG2 and HG4, at 1x MIC concentration had increased survival rate by ~20% (Fig.7b). In comparison to larvae in the untreated control group, larvae infected with MRSA USA300 LD, followed by treatment with peptides 3x MIC increased survival by 4.6-fold and 4.4-fold for HG2 and HG4 with a survival rate of 78% and 75%, respectively (Fig. 7b).

We were able to show that both peptides, especially at 3x MIC, concentration can effectively control MRSA USA300 infection *in vivo* in *G. mellonella*. Other pharmacological aspects of the peptides need to be investigated in order to improve the efficacy of HG2 and HG4, as survival rate of AMP-treated larvae was comparable to the LD_50_ survival rate, probability due to distribution and/or bioavailability of the peptides *in vivo* (in *G. mellonella* model). Nonetheless, our results show that the peptides HG2 and HG4 at 3x MIC concentration are capable of significantly improving the survival of larvae infected with a lethal dose MRSA USA300 and are active against this clinically important drug resistant pathogen in an *in vivo* model.

## Conclusion

The two AMPs, HG2 and HG4, identified using *in silico* approaches from a rumen metagenomic dataset have further confirms the rumen as an invaluable resource for urgently needed alternatives to currently available antibiotics. Furthermore, study presented here emphasises the usefulness of complementing wet-lab and *in silico* techniques for the rapid identification of new AMP candidates from environmental samples. The low similarity of the newly identified AMPs to previously known sequences suggest their novelty from an evolutionary point of view. Experimental evaluation and characterisation of the antimicrobial properties of HG2 and HG4, two of the identified AMP candidates revealed their antimicrobial activity against Gram positive bacteria. Their activity against MRSA suggest that membrane permeabilisation and decrease in intracellular ATP concentration might play a role in their antimicrobial activity. HG2 and HG4 both preferentially bind to MRSA total lipids rather than with human erythrocyte lipids. HG4 was less cytotoxic against all cell lines tested and was observed to bind more specifically to pure bacterial membrane lipids, indicating that HG4 may form a more superior template for a safer therapeutic candidate than HG2. The non-toxic effect of the peptides against *G. mellonella* larvae, and their *in vivo* efficacy against MRSA USA300 infection in the *G. mellonella* infection model suggests that these peptides might possess potential as safe alternative therapeutic agents with anti-biofilm activity for the treatment of bacterial infections. Given the technological advances, improvements in genomic methods and computational analytic approaches as well as the growing abundance of omics data, it is likely that the approach developed and presented here for the identification of novel AMPs, might facilitate the discovery of a growing number of other AMPs and other bioactives from environments where conventional isolation and cultivation of microorganisms is challenging.

## Materials and Methods

### *In silico* prediction and identification of AMPs using computational analysis

Antimicrobial peptide prediction and similarity analysis was performed on the rumen metagenomic dataset from the study by Hess et al,^19^. The dataset was termed the Library “Cow” dataset and contains 2,547,270 predicted protein sequences (‘metagenemark_predictions.faa.gz’) which were downloaded from the weblink (http://portal.nersc.gov/project/jgimg/CowRumenRawData/submission/). All other datasets (libraries) used for similarity analysis prediction/identification of novel AMP candidates from the “Cow” dataset and their respective sources are as follows: The Library “AMP1”, which contained a list of 2308 known antimicrobial peptides (AMPs) downloaded from APD2^34^ (downloaded on November 10, 2013 and available at http://aps.unmc.edu/AP/main.php) and the Library “AMP2”, which contains of a list of 48 synthetic AMPs (Hilpert Library) identified by Ramon-Garcia et al^35^. The MATLAB toolbox Gait-CAD and its successor SciXMiner (http://sourceforge.net/projects/scixminer/)^86, 87^ including the Peptides Extension Package^88^ were mostly used for the computational analysis.

The “fastread” function of MATLAB Bioinformatics toolbox was used to import the “Cow” dataset. For easier computational analysis, the imported dataset was then split into 26 parts with ~100,000 sequences each. Following recommendations that small antimicrobial proteins should have a length <200 amino acids (AAs), with most AMPs (>90%) on the APD2 database having a length of <60 AAs^34^, only protein sequences with a maximum length of 200 AAs^33^ and not more than 5% unknown AAs (marked by X, *) were selected from the “Cow” dataset predicted protein sequences (metagenemark_predictions.faa.gz). Libraries “AMP1” and “AMP2” were combined to produce Library “AMP” composing of a total of 2356 peptides. Thereafter, AA distribution and AA dimer (pair) distribution were computed, resulting in proportion for 20 AAs (and 20×20 = 400 AA dimers) for Libraries “Cow” and “AMP”. Pairwise distances of AA acid distributions between two peptides (termed “AAD”) were computed with a minimal value of 0 for identical and increasing values for different AAs. Pairwise distances of AA acid and AA acid pair distributions between two peptides were computed respectively (distance for 400 + 20 features), termed “AAPD”. For each candidate of Library “Cow”, the values of “AAD” and “AAPD” to each peptide in Library “AMP” were computed. Again, for each sequence in Library “Cow”, minimal distance values from the previous computational step and the number of most similar peptide from Library “AMP” were saved as separate features. To select only promising candidates of AMP predictions, only sequences from the Library “Cow” with small AA distances AAD <0.2 or a small AA and AA pair distances AAPD <1.45 were saved. The distances were computed using the 1-norm (Manhattan norm). The thresholds for AAD and AAPD were heuristically chosen to balance the trade-off between too many weak candidates (too high values of AAD and AAPD) vs. the loss of promising candidates (too low values of AAD and AAPD). All sequences that fulfilled the conditions in the preceding step were collected and some interesting hits were used to check similarity in APD2 database (criteria: small values of AAD or AAPD, different neighbours to explore the variety of the candidates found, short peptide length). Finally, descriptors were computed following procedures described by Mikut et al^88, 89^ to check the expected balance between hydrophobicity and positively charged AAs as a typical design criterion for AMPs.

### Peptide synthesis and 3-dimensional molecular modelling of peptide structures

Pure peptides (≥95% purity) were synthesized on resin using solid phase Fmoc peptide chemistry^39^ by GenScript Inc. USA. A *de novo* structural prediction method, PEP-FOLD^49^ was used to model the 3D conformation of peptides HG2 and HG4. 200 simulations were ran for each peptide and resulting conformations where clustered and ranked using the sOPEP coarse grained force field^90^. In the case of HG2 the formation of the cysteine bond between Cys10 and Cys18 was imposed as restraint to the simulation. The best 3D models for each peptide was chosen and manually analysed using PyMOL v1.7.6^91^.

### Antimicrobial susceptibility of bacterial cells

To determine the antimicrobial activity of new antimicrobial peptides, HG2 and HG4, their minimum inhibitory concentrations (MICs) were determined by broth microdilution method^92^ in cation adjusted Mueller Hinton broth (MHB) following the International Organization for Standardization (ISO) 20776-1 standard for MIC testing using a final bacterial inoculum of 5 × 10^5^ CFU/ml^93^. The lowest concentration of the AMPs that inhibited the visible growth of the bacteria tested after an overnight incubation at the appropriate temperatures (37°C for all organisms except for *Listeria monocytogenes* 30°C) and growth conditions was taken as the MIC. The peptides dissolved in sterile distilled water were added to bacteria culture and incubated overnight at appropriate conditions. The MICs of the peptides and comparator antibiotics were recorded after 18-24 h. The minimum bactericidal concentrations (MBCs) were also determined and were taken as the lowest concentration of the antimicrobial that prevented the growth of bacterial cells after subculture of cells (from MIC treatment) onto antibiotic-free media.

### Time kill kinetics

The bactericidal activity of HG2, HG4 and comparator antimicrobial compounds was assessed as previously described^94^ using exponential-phase cultures of MRSA USA300 grown in MHB (1 × 10^6–8^ CFU/ml). Cells were treated with antimicrobial compounds at 3 times their MIC concentrations (final concentrations), incubated 37°C with gentle shaking at 110 rpm. Samples were taken at different time points and inoculated onto MH agar plates using the spread plate technique. After overnight incubation, the colony forming units per millilitre of cell culture (CFU/ml) was calculated. Experiments were performed in quadruplicates.

### Anti-biofilm activity

The ability of HG2 and HG4 to disrupt established *S. aureus* biofilms was measured using a 96 well format as described by^15^. MRSA USA300 cultures grown overnight in Brain Heart Infusion (BHI) broth was re-suspended to an OD_600nm_ = 0.02 and grown without shaking at 37°C in 96 well tissue culture plates for another 24 h. The planktonic cells were removed by three PBS (phosphate buffered saline) washes. Thereafter, fresh BHI broth containing peptides HG2 or HG4 at sub- and supra MIC concentrations (0.5X, 1X, 2X and 4X MIC) was added to wells containing adherent cells and incubated without shaking for another 24 h. Planktonic cells were again removed by three PBS washes. Biofilms were fixed with methanol for 20 min, stained with 0.4% (w/v) crystal violet solution for 20 min and re-solubilised with 33% (v/v) acetic acid. The optical density of re-solubilised biofilms was measured at 570_nm_ in a microplate spectrophotometer. The growth of biofilm per treatment was calculated as a percentage of the untreated cells and anti-biofilm activity was determined for statistically significant treatments.

### Serial passage/resistance assay

*In vitro* evaluation of the potential for MRSA USA3000 cells to develop resistance to HG2 and HG4 was performed as previously described^95^. Briefly, on Day 1, overnight cultures grown from a single colony of MRSA USA300 strain in MHB at 37°C with shaking at 225 rpm was subjected to microbroth dilution susceptibility testing performed using a standard doubling-dilution series of AMP concentrations as described for MIC determination. Cultures from the highest concentration that supported growth were diluted 1:1000 in MHB and used to provide the inoculum for the next passage day. This process was continued for 20 days. Any putative mutants recovered were colony purified for three generations on MHA, prior to further characterization.

### ATP determination assay

Adenosine triphosphate (ATP) drives many cellular and metabolic processes and can be used to ascertain the integrity of cells^75, 76^. To determine whether HG2 and HG4 affected ATP levels in *S. aureus* treated cells, we used the ATP colorimetric/fluorometric assay kit (Sigma Aldrich) which determines ATP concentration by phosphorylating glycerol, resulting in a colorimetric product that shows absorbance at 570nm. Briefly, in a clear 96 well plate, samples and ATP standard provided with the kit were added in a reaction mix containing ATP assay buffer, ATP Probe and Converter and Developer mix. The reaction mix was incubated in the dark at room temperature for 30 min. The absorbance at 570nm was then measured in a microplate spectrophotometer. The background ATP levels obtained from samples and ATP standards were autocorrected by subtracting ATP levels from blank treatments. The amount of ATP in unknown samples were determined from the ATP standard curve and calculated using the formula in the instruction manual. All samples were tested in triplicates.

### Membrane permeabilisation assay

Permeabilisation of the bacterial cytoplasmic membrane by HG2 and HG4 peptide was evaluated using the cell-impermeable DNA probe propidium iodide as previously described^15, 78^. Logarithmic phase bacterial suspension of MRSA USA300 was prepared by diluting overnight cultures in fresh MHA broth (1 in 10 dilution), incubated 3 h at 37°C, 200 rpm, and pelleted by centrifugation for 5 min at 3000 g. Bacterial cell pellet was then resuspended in sterile PBS at about 10^9^ bacteria/ml. Propidium iodide (at 1 mg/ml, Sigma Aldrich) was added to the bacterial suspension at a final concentration of 60 μM. This suspension (100 μl) was then transferred into black 96-well plates already containing 100 μl of serially diluted HG2 or HG4 peptide in PBS. Kinetics of fluorescence variations (excitation at 530nm / emission at 590 nm) were then recorded using a microplate reader over 80 min period with incubation at 37°C. Cetyl trimethylammonium bromide (CTAB) (at 300 μM) served as positive control giving 100% permeabilisation. The permeabilisation effect of HG2 and HG4 were expressed as percentage of total permeabilisation.

### Transmission electron microscopy

Transmission electron microscopy was used to investigate the effects of HG2 and HG4 on *S. aureus* cell morphology as described by^15^. Mid-log phase *S. aureus* cultures treated with HG2 and HG4 (at 3× MIC for 1 h) were fixed with 2.5% (v/v) glutaraldehyde and post-fixed with 1% osmium tetroxide (w/v). They were then stained with 2% (w/v) uranyl acetate and Reynold’s lead citrate after which they were observed using a JEOL JEM1010 transmission electron microscope (JEOL Ltd, Tokyo, Japan) at 80 kV.

### Haemolytic activity

The ability of HG2 and HG4 to cause leakage of erythrocytes from human whole red blood cells was determined to ascertain probable cytotoxicity to mammalian cells and the suitability of peptides for use as therapeutic agents. The haemolytic activity of HG2 and HG4 was determined as previously described^15^. Briefly, fresh human erythrocytes (obtained from Divbio Science Europe, NL) were washed 3 times by centrifugation at 800 g for 5 min with sterile phosphate buffer saline (PBS, pH 7.4). The washed erythrocytes were resuspended in PBS to a final concentration of 8%. 100 μl of human erythrocytes were then added per well into sterile 96 well microplate already containing serial dilutions of the peptides in 100 μl of PBS. The treated red blood cells were incubated at 37°C for 1 h and centrifuged at 800 g for 5 min. The supernatant (100 μl) were carefully transferred to a new 96 well microplate and absorbance OD_450nm_ measured using microplate reader. Triton-X100 at 0.1% (v/v) was used as positive control giving 100% haemolysis and haemolysis caused by HG2 and HG4 was expressed as percentage of total haemolysis. The HC_50_ values for HG2 and HG4 (i.e. the concentration of peptide causing either 50% of haemolysis) were calculated using GraphPad® Prism 7 software.

### Peptide-lipid interaction and insertion assay

Peptide-lipid interaction was measured using lipid monolayer formed at the air:water interface with total lipid extracts and pure lipids. Total lipids were extracted from overnight cultures of MRSA or human erythrocytes using Folch extraction procedure as previously described^15, 78, 96^. Extracted total lipids were dried, resolubilised in chloroform:methanol (2:1, v/v) and stored at −20 °C under nitrogen. Pure prokaryotic and eukaryotic lipids used were: cardiolipin, POPC (1-palmitoyl-2-oleoyl-glycero-3-phosphocholine), POPE (1-palmitoyl-2-oleoyl-sn-glycero-3-phosphoethanolamine) and POPG (1-palmitoyl-2-oleoyl-sn-glycero-3-phospho-(1’-rac-glycerol) (Avanti Polar Lipid). LTA (Lipoteichoic acid from *S. aureus*) and LPS (lipopolysaccharide from *E. coli*) (both obtained from Invitrogen) were also tested. Pure lipids were reconstituted in chloroform:methanol (2:1, v/v) at 1 mg/ml and stored at −20°C under nitrogen. For peptide-lipid interaction assay, lipid monolayers at the air:water interface were formed by spreading total lipid extract or pure lipids at the surface of 800 μl of sterile PBS using a 50 μl Hamilton’s syringe. Lipids were added until the surface pressure reached the desired value. After 5-10min of incubation allowing the evaporation of the solvent and stabilization of the initial surface pressure, 8 μl of HG2 or HG4 diluted in sterile PBS at 100 μg/ml were injected into the 800 μl sub-phase of PBS under the lipid monolayer (pH 7.4, volume 800 μl) using a 10 μl Hamilton’s syringe giving a final concentration of peptide of 1 μg/ml, as preliminary experiments had shown that this concentration was optimal. The variation of the surface pressure caused by peptide insertion was then continuously monitored using a fully automated microtensiometer (μTROUGH SX, Kibron Inc) until it reached equilibrium (maximal surface pressure increase being usually obtained after 15-25min). To reflect physiological situations, the initial surface pressure was fixed at 30 ± 0.5 mN/m in some experiments as this value corresponds to a lipid packing density theoretically equivalent to that of the outer leaflet of the eukaryotic and prokaryotic cell membrane^97^ In other experiments, the critical pressure of insertion of HG2 or HG4 in the total lipid extracts and pure lipids was measured as previously described^15, 96^. Briefly, in these experiments, the initial pressure of lipid monolayer was set-up at different values (between 10 and 30 mN/m) and the variation of pressure caused by the injection of peptide was measured. Critical pressure of insertion was calculated by plotting the variation of surface pressure caused by peptide insertion as a function of the initial surface pressure, and corresponds to the theoretical value of initial pressure of lipid monolayer that does not allow the insertion of the peptide, i.e. a variation of pressure equal to 0 mN/m. All experiments were carried out in a controlled atmosphere at 20 °C ± 1 °C and data were analyzed using the Filmware 2.5 program (Kibron Inc.). The accuracy of the system under our experimental conditions was determined to be ± 0.25 mN/m for surface pressure measurements.

### Human cell culture and cytotoxicity studies

The toxicity of HG2 and HG4 was tested as previously described^98–100^. The following human cells were used: BEAS-2B (normal airway epithelial cells, ATCC CRL-9609), IMR-90 (normal fibroblasts, ATCC CCL-186) and HepG2 (liver cell line, ATCC HB-8065). BEAS-2B, IMR-90 and HepG2 cells were cultured in Dulbecco’s modified essential medium (DMEM) supplemented with 10% fetal calf serum (FCS), 1% L-glutamine and 1% antibiotics (all from Invitrogen). Cells were routinely grown onto 25 cm^2^ flasks maintained in a 5% CO_2_ incubator at 37°C. Briefly, cells grown on 25 cm^2^ flasks were detached using trypsin-EDTA solution (Thermofisher) and seeded into 96-well cell culture plates (Greiner Bio-one) at approximately 10^4^ cells per well (counted using Mallasez’s chamber). The cells were grown at 37°C in a 5% CO2 incubator until they reached confluence (approximately 48-72 h of seeding). Wells were then aspirated and increasing concentrations of HG2 or HG4 were added to the cells and incubated for a further 48 h at 37°C in a 5% CO_2_ incubator. The wells were then emptied, and cell viability was evaluated using resazurin based *in vitro* toxicity assay kit (Sigma-Aldrich) following manufacturer’s instructions. Briefly, resazurin stock solution was diluted 1:100 in sterile PBS containing calcium and magnesium (PBS++, pH 7.4) and emptied wells were filled with 100 μl of the resazurin diluted solution. After 4 h incubation at 37°C with the peptide treated cells, fluorescence intensity was measured using microplate reader (excitation wavelength of 530 nm/emission wavelength of 590 nm). The fluorescence values were normalized by the controls and expressed as percentage of cell viability. The IC_50_ values of HG2 or HG4 on cell viability (i.e. the concentration of peptides causing a reduction of 50% of the cell viability) were calculated using GraphPad® Prism 7 software.

### *In vivo* efficacy in *Galleria* model

All procedures of larvae rearing, injection and *G. mellonella* killing assays were conducted as previously described^101^. In each assay, ten (10) larvae weighing between 280 - 300 mg each were randomly selected. Larvae with previous melanisation of the cuticle were not used in the experiments. All the experiments were designed in at least four experimental and biologic replicates.

Firstly, we evaluated the putative toxic effect of the peptides HG2 and HG4 in *G. mellonella*. Each peptide solution prepared in sterile H_2_O was injected in larvae at the concentration of 1x MIC, as 16 mg/kg of larvae body weight (LBW) for HG2 and 32 mg/kg LBW for HG4; and 3x MIC 64 mg/kg LBW HG2 and 98 mg/kg LBW HG4 as previously determined. After injection with peptides, the larvae were maintained at 37°C in the dark. Phenotypic aspects such as melanisation and mobility of the larvae as well as survival were monitored every 24 hours for 96 hours. Larvae not inoculated with APMs were used as controls.

To determine the LD50 and LD of *S. aureus* MRSA USA300 in *G. mellonella*, an inoculum of 10 μl of MRSA USA300 suspension in PBS 1X (10^3^ to 10^6^ CFU/larva) was injected into the larvae hemocoel using insulin syringes (Becton Dickinson, USA). Larvae inoculated with PBS and larvae not inoculated were used as negative controls. After the injections, the larvae were maintained at 37°C in the dark, the survival was recorded every 24 hours for 96 hours and the LD50 and LD were determined.

The efficacy of the peptides HG2 and HG4 to control MRSA USA300 infection *in vivo*, was evaluated following the protocol by Peleg et al.,^84^ The groups of *G. mellonella* larvae were infected with the predetermined lethal dose of MRSA USA300. After 30 minutes of the larvae injection with the bacteria, the peptides were injected at 1x MIC and 3x MIC concentrations respectively. Larvae injected with PBS solution, and HG2 or HG4 peptides solution (1x MIC or 3x MIC) alone were used as negative controls. Injected larvae were maintained at 37°C in the dark and their survival was monitored and analyzed as above.

### Statistical analysis

All biological experiments were repeated at least three times and three biological replicates were used wherever applicable. Results are expressed as mean ± standard error. The MRSA USA300 LD50 value was calculated by linear regression using software R v.2.13.0^102^. The Kaplan-Meier method was used to plot the survival curves. Differences in survival were calculated using the log-rank test with the software SigmaPlot from Systat Software Inc., San Jose, California^103^. A *P*-value of 0.05 was considered to be statistically significant.

### Author Contributions

LO and SH, and CC conceived the project. LO, with help from HV, TW and MW, completed the laboratory work under supervision of SH and CC. AC and NF assisted LO with transmission electron microscopy and 3D structural modelling respectively. LO, HO, MM and JP completed the membrane permeabilisation, cytotoxicity and lipid interaction assays. LO, and JAP completed the ATP assays. KH and RM assisted LO with screening of metagenomic library and identification of AMP candidates. AT, MG completed *in vivo* work in *G. mellonella* with input from LO and supervision by DB, HM and SH. MH and HM have provided valuable ideas into the project from the time of conception. LO wrote the paper with input from all co-authors.

## Acknowledgement

This project was funded partly by the Cross River State Government of Nigeria, the Life Sciences Research Network Wales, RCUK Newton Institutional Link Fund (172629373), and the BBSRC UK (BB/L026716/1). HM thanks the Coordination for the Improvement of Higher Education Personnel (CAPES) for providing the Joint Institutional Links grant. KH thanks the Institute of Infection and Immunity of St. George’s University of London for a start-up grant. We are also grateful to Dr Colin Greengrass, for his advice. The authors declare no competing financial or other interests.

## Competing Interests

The authors declare no competing interests.

## Supporting Information Legends

**Fig. S1. Mass spectrometry and certificate of analysis (COA) report for AMP chemical synthesis:** a) HG2 and b) HG4. Peptides were synthesised by GenScript Inc. USA. Mass spectrum analysis and COA report show correct molecular weight and purity grade for both peptides.

**Table. S1 Summary of results from computation steps**

**Table. S2 Six most promising antimicrobial peptide candidates (HG1-HG6) identified in cow rumen metagenome dataset**

**Table. S3 Scaffold nucleotide sequences from which putative antimicrobials H-G1-H-G6 were identified**

**Table. S4 Peptide lipid interaction and insertion measurements:** interaction of peptides, HG2 and HG4 with total MRSA and erythrocyte lipid extracts, and interaction of HG2 and HG4 (at 1 *μ*g/mL final concentration) with pure lipids the initial surface pressure of lipid monolayer. Maximal variation of surface pressure induced by the injection of peptide in lipid monolayer with initial surface pressure of 30+/−0.5 mN/m.

